# Alanine catabolism of Paneth cells maintains intestinal stem cell function during dietary restriction

**DOI:** 10.1101/2025.08.28.672976

**Authors:** Ashish Kumar, Sharif Iqbal, Ernesta Nestaite, Nalle Pentinmikko, Martin Broberg, Jack Morikka, Ville Hietakangas, Pekka Katajisto

## Abstract

**Background and aims:** Paneth cell niche promotes the function of intestinal stem cells (ISCs) during reduced food intake, but how ISC activity can be boosted when availability of resources is simultaneously reduced remains unknown.

**Methods:** Mice were subjected to dietary restriction (DR) and *ad libitum* (AL) dietary regime. FACS-sorted DR and AL Paneth cells and Lgr5+ stem cells were co-cultured, and organoid formation supportive capacity of Paneth cells was assessed. The role of Gpt2 – a Paneth cell specific mitochondrial alanine transaminase, was investigated by targeting pharmacologically and genetically in the intestinal organoids. By using U-^13^C labelled Alanine, an intercellular metabolic exchange assay was developed – and a conversion and transport of lactate from Paneth cells to stem cells, was followed by metabolomics. The *in vivo* role of *Gpt2* was investigated in a conditional knock out mouse model.

**Results:** We discovered that an increase in ISC function upon dietary restriction (DR) or DR mimicking condition, is dependent on Gpt2 in Paneth cells. Metabolic tracing of alanine directly showed DR boosts Paneth cell alanine catabolism, increasing lactate production via gluconeogenesis. Alanine derived lactate is shuttled from Paneth cells to neighboring stem cells, promoting TCA cycle and enhancing the ISC function during DR. Correspondingly, pharmacologic inhibition i*n vitro* and genetic targeting of *Gpt2 in vivo* abolished the DR induced capacity of Paneth cells to support ISC function.

**Conclusions:** Our results unravel dynamic intercellular metabolic cooperation in tissue adaptation, where Paneth cells upregulated Gpt2 catabolize increased amount of alanine to generate lactate as a transferable energy source, and to augment stem cell functions under DR.

## Introduction

Reduced intrinsic nutrient status robustly inhibits cell proliferation and organ growth ^1^. However, the number and function of actively cycling intestinal stem cells increases during DR ^2^. How tissue resident stem cells can circumvent the growth inhibitory effects of low energy state is poorly understood.

The intestinal epithelium is a rapidly renewing tissue responsible for food absorption and the barrier between the host and the microbiome. The rapid renewal is maintained by actively cycling intestinal stem cells (ISCs), which reside in the crypts of Lieberkühn and express *Lgr5* ^3^. ISCs are intermingled with Paneth cells, which constitute a key component of the ISC niche by providing factors regulating ISC self-renewal and differentiation ^4^. Interestingly, alterations in either nutrient quality or quantity have recently been shown to alter stem cell function. ISCs were shown to respond to high-fat ketogenic diet by Hmgcs2 mediated generation of beta-hydroxybutyrate (β-OHB), and to complete fasting by Cpt1a mediated fatty acid oxidation ^5, 6^. Additionally, moderate reduction in food intake over a longer period of time – referred to as Dietary Restriction (DR) – impacts stem cells in addition to intrinsic mechanisms ^7^ via the Paneth cell niche ^2^. During DR, the activity of the central integrator of cues on nutrient availability, mTORC1, is reduced and results in increased expression of *Bst1* in Paneth cells ^2^. This increases the production of extracellular cADPR, which promotes clonogenicity and the number of stem cells, while the proliferation of differentiating progenitors called Transiently Amplifying (TA) cells is reduced. As a result, DR reduces the generation of differentiated cells and the absorptive surface of the villi, but the stem cell population in the crypt is increased to allow rapid villus expansion if more food becomes available.

The exchange of metabolites between stem cells and their niche was recently described in selected tissues ^8, 9^. In the intestine, the cycling ISCs have an energy metabolism that is heavily reliant on oxidative phosphorylation (OXPHOS) ^8^. Paneth cells, on the other hand, produce high levels of lactate as a result of glycolysis. Interestingly, the lactate produced by Paneth cells can be exported and thereafter could provide fuel for the neighboring ISCs to promote the stem cell functions ^8^. However, whether the lactate used by stem cells to fuel their mitochondrial OXPHOS actually originates from Paneth cells has not been directly demonstrated. Moreover, other works show that inhibition of pyruvate entry into the mitochondria in ISCs increases their self-renewal and function ^10^. Consequently, the relevance of metabolic interplay between cell types, and the role of mitochondrial metabolism in determining ISC function remain obscure.

As reduction in nutrient intake reduces the production of differentiated intestinal cells by progenitors, but concomitantly the number and function of ISCs increases, we asked whether the exchange of metabolites between stem cells and their Paneth cell niche provides means to maintain stem cell function during nutritional challenges. We found that during DR, Paneth cells metabolize their own alanine pool to generate lactate that serves as a metabolic currency to be transferred to the stem cells. The alanine catabolism discovered here presents the first dynamically regulated metabolite interaction between stem cells and the niche. Our results highlight the importance of niche interactions, and how response of stem cells can differ from progenitors lacking the metabolic support of the niche.

## Materials and Methods

### Animals

*Lgr5-eGFP-creERT2* mice were maintained in a C57BL/6J background (Barker *et al*., 2007). Exon 4 of Gpt2 was flanked with LoxP sites to create *Gpt2* ^*fl/fl*^ mice and crossed with *Lgr5-eGFP-creERT2* to generate *Lgr5-eGFP-creERT2;Gpt2* ^*fl/fl*^ mice. Mice (2-3 mo) were then fed with tamoxifen (TAM) chow (Envigo) for 2 weeks. TAM-fed mice (*Gpt2*^*WT*^ and *Gpt2*^*cKO*^) were maintained for 90 days under AL diet, and subsequently subjected to AL or DR (30% reduction of food intake compared to their AL counterparts) in pairwise manner for 15 days before analysing. Rapamycin (4mg/kg) was injected intraperitoneally for consecutive 10 days as demonstrated previously^11^. Animal experimentations were approved by the Finnish National Animal Experiment Board.

### Media

ENR medium was used to culture isolated crypts and single cells (media composition, Table S2).

ENR mimetic medium: Minimum essential media (2109022, Gibco) composition was compared with DMEM F/12 Advanced (Gibco) composition to formulate ENR mimetic media and required components were added (Table S1). Alanine is deficient in EMM media and added according to the experimental requirements.

### Immunoblotting

Western blot analysis has been done as earlier^11^. Antibodies details are in Table S2.

### Immunohistochemistry

Immunohistochemistry was performed as previously^11^. Briefly, tissues were fixed in 4% PFA, paraffin-embedded and sectioned. For antigen retrieval boiled 1X citrate buffer (Sigma-Aldrich), pH 6.0 was used for 15 min. Subsequently, permeabilization was performed with 0.5% Triton-X100 (Sigma). Sections were incubated with primary antibodies (Table S2) overnight at 4^º^C and detected with biotin-conjugated secondary antibodies and DAB substrate on a peroxidase-based system (Vectastain Elite ABC, Vector Labs).

### Immunofluorescence staining of isolated crypts

Isolated crypts were fixed with 4% PFA for 15min RT and washed once with 0.2% BSA-PBS. Crypts were permeabilized with 0.5% Triton X100 and incubated in blocking solution (5% Normal Goat Serum - 0.2% Triton X-100 - 0.1%BSA PBS). Crypts were stained with primary antibodies (Table S2) at 4^º^C, washed and incubated with secondary antibodies (2h, RT). Crypts were washed, followed by nuclear co-stain (Hoechst 33342) for 30 min before imaging.

### Real-time qPCR

RNA from isolated crypts, cultured organoids, and sorted Lgr5^hi^ and Paneth cells was isolated by Trizol purification (Life). Isolated RNA was transcribed with cDNA synthesis kit (Molecular probes). qPCR amplification was detected using SYBRGreen (2xSYBRGreen mix, Applied biosciences) method. Primers details, Table S2.

### Isolation of intestinal crypts and organoid culture

Intestinal crypts isolation was performed as mentioned previously^11^. Briefly, small intestines were flushed in cold PBS and cut into pieces. Crypts were dissociated by robust shaking in 10mM EDTA-PBS on ice for two hours, passed through 70 μM nylon mesh, washed and plated (~300 crypts per 25μl drop of 60% Matrigel (BD Biosciences)). Crypts were cultured with ENR medium and Y-27632 (10μM) was added for the initial 2 days. Starting frequency (Day-2) and regenerative potential (crypt domains per organoid, at day 5-6) of primary organoids were assessed.

### FACS based sorting of single cells and co-culture

The isolated crypts or grown organoids were dissociated in TrypLE express (Gibco) with 1000U/ml of Dnase I (Roche) at +32^º^C (90 sec for crypts, 5 min for organoids at +37^º^C). Dissociated cells were stained with the following antibodies (Table S2) and sorted as described previously^11^. Cells were sorted using FACSAria Fusion (BD Biosciences) with an 85μm nozzle. Sorting strategy: Intestinal stem cells, Lgr5-eGFP^hi^Epcam^+^CD24^med^SYTOXblue^−ve^ CD31^−ve^Ter119^−ve^CD45^−^ve-SYTOXblue^−ve^ and Paneth cells, CD24^hi^ SSC^hi^ Epcam^+ve^CD31^−ve^Ter119^−ve^CD45^−^ve-SYTOXblue^−ve^. Post sorting, equal numbers of Lgr5^hi^ and Paneth cells were added to Matrigel and cultured with ENR media supplemented with 1 μg/ml R-spondin-1, 100 ng/ml Wnt3A (RnD), 1 μM Jagged-1 peptide (Anaspec). Single cell-starting frequency and colony formation potential of primary organoids (day 6) and budding per organoid (day 10) was scored.

### Metabolomics

For metabolomic samples, the isolated crypts were dissociated and ISCs and Paneth cells were FACS sorted. The sorted cells were washed with ice-cold PBS and resuspended in metabolomics extraction buffer (Acetonitrile/dH2O; 80:20). For tracing experiments with isotope labelled alanine, isolated crypts from AL and DR mice were incubated with ENR mimetic media (lacking alanine) supplemented with 4mM ^U13^C3-Alanine (Cambridge isotope laboratories) for 45 mins at +37^º^C. Crypts were dissociated and the single cell suspension was prepared using TrypLE express (Gibco) with 1000U/ml of Dnase I (Roche) at +32^º^C for 90 sec. Paneth and intestinal stem cells were sorted, washed and resuspended in metabolomics buffer. The metabolites peak areas in sorted Paneth and Lgr5^hi^ cells were calculated and analyzed as previously ^12^ and values were normalized to the cell numbers (Paneth and Lgr5^hi^) per condition and represented as fold change in ratio of DR to AL.

### CRISPR-Cas9 editing of *Gpt2* in organoids and rapamycin treatment

#### Plasmid construction for CRISPR guides, lentiviral production and transduction

Guides were designed for targeting *Gpt2 (exon 2)* using the CRISPR design tool (https://portals.broadinstitute.org/gppx/crispick/public) (Table S2). Designed oligos were annealed, and cloned into the pLentiCRISPRv2 plasmid. Using Lipofectamine 2000 (Invitrogen), lentiviruses were produced by co-transfecting transfer vectors (pLentiCRISPRv2 gGpt2, and pLentiCRISPRv2 gScramble) and packaging vectors (CMV-VSVg and Delta8.9) into 293FT cells cultured in DMEM (Sigma-Aldrich). After 40 hours of transfection, collected viral particles were concentrated 100x using a Lenti-XTM concentrator (Clontech). Transduced organoids were selected for 3 days (2 μg/ml puromycin)^13^.

Knockout efficiency of *Gpt2*^*KO*^ organoids was confirmed by sequencing, RT-PCR and immunoblotting. To mimic DR effect *in-vitro*, Wild-type and *Gpt2*^*KO*^ organoids were seeded for initial 2 days without treatment and 2nM Rapamycin (Selleckchem) was supplemented for next 2 days, and repeated this cycle for 16 days. Then, Paneth cells were FACS sorted for Rapamycin primed organoids. Co-culture was established with sorted Lgr5^hi^ stem cells isolated from AL mice.

### Statistical Analysis

Minimum of *n* = 3 independent experiments were performed unless otherwise mentioned, and the numbers of replicates are indicated in the figure legends. Investigators used blind analysis to analyze *in-vitro* organoid cultures, but it was not always possible for co-culture experiments due to its nature. Graphpad Prism v.9.0.0 was used for statistical analysis and data plotting. An unpaired two-tailed Student’s t-test was used to analyze the data; exact p-values are presented in the corresponding figures. p ≤ 0.05 is considered significant.

## Results

### Niche-specific induction of Gpt2 boosts the regenerative potential of ISCs during DR

Reduction in nutrient intake boosts the regenerative potential of intestinal stem cells both intrinsically ^5, 7^ and via the niche ^2^, but the associated metabolic crosstalk between the intestinal stem cells and Paneth cell niche during DR remains poorly characterized. To explore this, we mined our previously published RNA sequencing data ^11^ for metabolism-related genes with cell type-specific expression patterns between Paneth cells and Lgr5^hi^ ISCs. Since the central integrator of amino acid sensing, mTORC1, and its downstream signaling have been implicated as the key regulator of nutrient-dependent responses in the intestine ^2, 5, 7^, we focused on genes related to amino acid metabolism. We found that a large subset of genes responsible for amino acid metabolism are differentially expressed between Paneth cells and ISCs (Fig.1A), and selected candidates based on consistent expression patterns in other previously published single-cell RNA sequencing data ^14^. One of the most interesting candidate genes was the mitochondrial alanine aminotransferase *Gpt2*, found to be expressed almost exclusively in Paneth cells (Fig. 1A). Gpt2 catalyzes the reversible conversion of alanine to pyruvate, which can be further converted to support various cellular functions ^15, 16^. To investigate the effect of dietary restriction on Gpt2, we subjected mice to *ad-libitum* or 30% DR diets for 15 days, and found that Gpt2 was specifically increased in the Paneth cells during DR (Fig. 1B and C, Fig. S1A). Of note, DR reduced mTORC1 activity in Paneth cells which coincided with an increase in ISC and Paneth cell numbers (Fig. S1B), like previously reported ^2^.

**Fig. 1:**
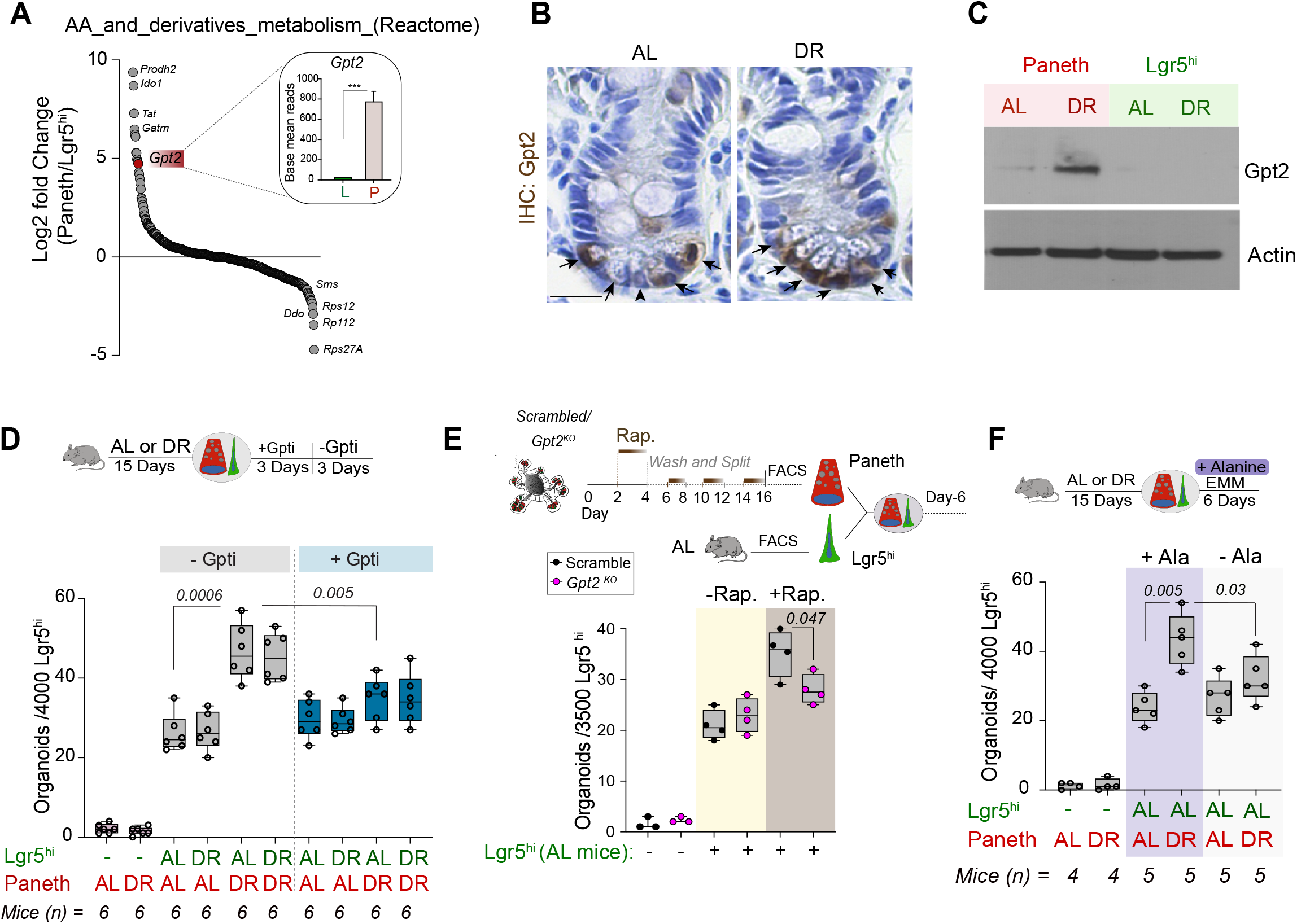
Increased Gpt2 activity in Paneth cells regulates ISC function under dietary restriction. **(A)** Base mean reads of genes involved with “Metabolism of amino acids and derivatives” (Reactome) (Jassal *et al*, 2020; Vastrik *et al*, 2007). Paneth cells (P) compared to Lgr5^hi^ (L) cells. *Gpt2* is expressed at a significantly higher level in Paneth cells (*p*=3×10^−37^). Differential expression analysis and statistical tests were performed using the DESeq2 method in R, obtained from the previously published data ^11^. **(B)** Immunohistochemistry for Gpt2 level in AL versus DR mice. The black arrow shows the Paneth cells. The image represents one of 4 biological replicates per condition. Scale Bar 20μm. **(C)** Immunoblotting of sorted Paneth and Lgr5^hi^ stem cells from AL and DR mice for Gpt2 shows an increase in DR Paneth cells (n = 3 mice per group). **(D)** Lgr5^hi^ cells (AL or DR) co-cultured with Paneth cells (AL or DR), treated with Gpt inhibitor (Gpti) for initial 3 days, and organoid formation potential of Lgr5^hi^ cells was calculated on day 6 (n = 6 mice). **(E)** Paneth cells were isolated from Scrambled and *Gpt2*^*KO*^ organoids which were treated with Rapamycin (2nM) for 8 days, with 2 days interval, after every 2 days of treatment, in total for 16 days and mixed with sorted Lgr5^hi^ cells isolated from AL mice. Clonogenic capacity of Lgr5^hi^ cells was calculated on day 6 (n=3-4 experimental replicates as indicated in figure). **(F)** Organoid formation potential of Lgr5^hi^ (AL or DR) co-cultured with Paneth cells (AL or DR) in EMM medium with or without alanine (0.5mM) was scored on day 6 (n=4-5 mice). Data are shown as mean ± s.d., and subjected to two-tailed unpaired Student’s *t*-test. *p* values shown in the corresponding panels. *p* <0.05 is considered statistically significant.

To investigate the possible functional relevance of Gpt2 in the niche mediated induction of stem cell function, we derived small intestinal organoids by co-culturing isolated Lgr5-Egfp^hi^ stem cells ^3^ and Paneth cells from AL and DR mice in all four combinations ^2^. Such co-cultures provide a strong tool for separately assaying the functional effects of DR on stem cells and their supporting Paneth cell niche. In line with our earlier findings ^2^, organoid formation by stem cells was significantly increased by DR treated Paneth cells, whereas ISCs showed no intrinsic response to DR (Fig. 1D). Interestingly, the enhanced capacity of DR Paneth cells to promote organoid formation was significantly reduced upon the addition of β-Chloro-L-alanine, an alanine aminotransferase inhibitor (Gpti) ^17, 18^ (Fig. 1D). Gpti can inhibit both the cytoplasmic Gpt1 and mitochondrial Gpt2. We found that Gpt1 is not expressed in the crypts where the Paneth cells and ISCs are located (Fig. S1C). Furthermore, organoid formation ability of AL Paneth cells (with low *Gpt2* expression) remained unaffected, further suggesting that the effect of Gpt inhibition was specific to the Gpt2. To validate our findings, we targeted Gpt2 in organoids using CRISPR-Cas9 and recapitulated the DR effects *in vitro* using mTORC1 inhibitor Rapamycin, a well-known DR mimetic (Fig. S1D and E). Similar to DR, Paneth cells isolated from Rapamycin treated organoids were able to enhance organoid formation capacity of AL Lgr5^hi^ stem cells (Fig. 1E). This was significantly reduced on Gpt2 knockout Paneth cells (Fig. 1E). Consistently, the rapamycin induced ability of Paneth cells to enhance the regenerative potential of AL stem cells was blunted by the Gpt inhibitor (Fig. S1F).

Dietary restriction is previously shown to increase the expression of the *Bst1* ectoenzyme in Paneth cells, resulting in increased production of cyclic-ADP-Ribose (cADPR) that promotes the function of the neighboring stem cells ^2^. Accordingly, cADPR increased the organoid forming capacity of co-cultures with AL Paneth cells (Fig. S2A). However, cADPR did not rescue the Gpti mediated suppression of organoid formation in co-cultures with DR Paneth cells (Fig. S2A-B). Taken together, Gpt2 activity in the Paneth cells promotes organoid formation by a mechanism that cannot be recapitulated with exogenous cADPR.

To validate the role of alanine in the DR-induced effects, we generated a customized organoid culture media (ENR mimetic media, EMM) lack alanine (Fig. S2C, Table S1). The DR induced boost in regenerative growth of crypts was significantly reduced in EMM (Fig. S2D), while EMM had no effect on crypts isolated from AL mice. Additionally, the alanine free culture medium suppressed organoid formation frequency only in crypts isolated from DR, not from AL mice (Fig. S2E). Finally, the boost in organoid formation by DR Paneth cells was significantly reduced in co-culture experiments lacking alanine (Fig. 1F). Together, these results indicate that the DR-induced Gpt2 in the Paneth cell niche promotes stem cell function in an alanine dependent manner.

### Alanine catabolism in Paneth cells generates lactate that can be taken by stem cells

Next, we wanted to understand the molecular nature of the support that Paneth cells provide to stem cells during DR. We developed a method for addressing alanine utilization in Paneth cells and stem cells, and the possible exchange of metabolites between the two cell types. Intestinal crypts we incubated immediately after their isolation with U-^13^C labeled alanine (m+3) for 45 minutes to specifically trace utilization of alanine and the metabolites derived from it. Freshly isolated crypts maintain their *in vivo* cellular organization and interactions between ISCs and Paneth cells (Fig. 2A). Consequently, the assay also allows exchange of metabolites between the cell types during the 45-minute labeling period, after which the crypts are rapidly disrupted into single cells and cell types are separated by cell sorting into the metabolomic analysis. We conducted alanine tracing on crypts isolated from AL or DR mice and noted that Alanine m+3 levels in DR Paneth cells were significantly increased in comparison to AL Paneth cells (Fig. 2B S3A). Alanine m+3 levels in ISCs did not change according to diet, indicating that alanine uptake is specifically increased by Paneth cells during DR. Consistently, previous RNA-seq data ^11, 14^ revealed that the expression of one of the alanine transporters, *Slc1A4*, is many folds higher in Paneth cells.

**Fig. 2:**
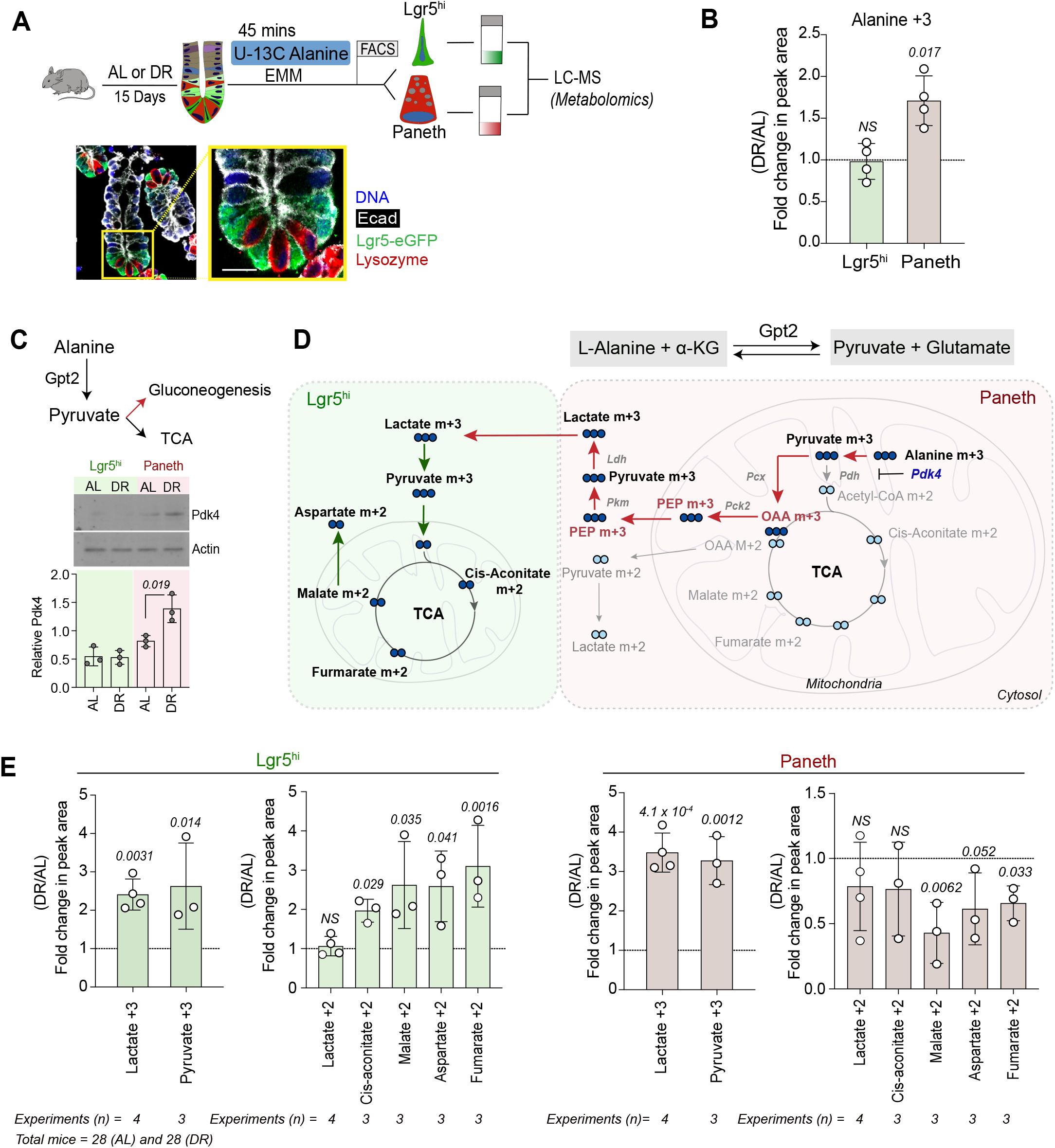
Alanine catabolism in DR Paneth cells forms lactate that supports TCA cycle in ISCs. **(A)** Schematic of the metabolomic tracing between two cell types, and a representative image of an isolated crypt with intact niche architecture. Nuclei (DAPI, blue), Paneth cells (Lysozyme, red), Stem cells (GFP, green), and E-cadherin (white). Scale Bar, 20μm. **(B)** Fold change in alanine m+3 in DR vs AL Lgr5^hi^ and Paneth cells. No. of experiments (n) = 4; Lgr5^hi^ and Paneth cells were sorted and pooled from 8-9 mice per condition (AL or DR) /experiment. Total number of mice: AL=28, DR= 28. **(C)** Immunoblotting for Pdk4 in sorted Paneth and Lgr5^hi^ stem cells from AL and DR mice indicates an increase in DR Paneth cells. Densitometric quantification (ratio to actin) (n = 3 mice per group). **(D)** Schematic illustrating possible metabolic routes of labelled alanine utilization in Paneth cells and mobilization of lactate to ISCs. **(E)** Fold change in different m+3 and m+2 labelled metabolites as indicated in DR versus AL in Lgr5^hi^ and Paneth cells. No. of experiments (n) = 3-4; Lgr5^hi^ and Paneth cells were sorted and pooled from 8-9 mice per condition (AL or DR) /experiment. Total number of mice: AL=28, DR= 28. Unless otherwise mentioned data are shown as mean ± s.d. with two-tailed unpaired Student’s *t*-test. *p* values shown in the corresponding panels. *p* <0.05 is considered statistically significant.

We also noted that DR increased the protein level of the Pdk4 kinase specifically in the Paneth cells (Fig. 2C). Pdk4 inhibits the pyruvate dehydrogenase (PDH) complex in mitochondria, and its high activity has been recently reported in less dividing intestinal epithelial cells ^19^. The increase Pdk4 in DR Paneth cells suggests that pyruvate may be shunted towards the gluconeogenic pathway instead of tricarboxylic acid cycle (TCA), similarly to gluconeogenic tissues like the liver ^20^.

The bidirectional reaction catalyzed by Gpt2 can generate pyruvate from alanine (Fig. 2D), which could provide a source of carbons under the DR conditions. Pyruvate utilization specifically in Paneth cells is reported to generate lactate ^8^, and on the other hand lactate is shown to fuel OXPHOS in ISCs. Interestingly, pyruvate m+3 and lactate m+3 were robustly increased both in Paneth cells and ISCs under DR, but lactate m+2 was not significantly altered in either of the cell types (Fig. 2E). As crypt cells do not express the cytoplasmic *Gpt1*, pyruvate m+3 is likely derived from alanine m+3 in the mitochondria by Gpt2.

Inside mitochondria pyruvate m+3 can either enter the TCA cycle where it will produce OAA (oxaloacetic acid) m+2 after loss of one heavy carbon, or it can be directly converted to OAA m+3 by pyruvate carboxylase (Pcx). OAA m+3 can convert to PEP m+3 (Phosphoenolpyruvate) in mitochondria by Pck2 to shuttle from mitochondria to cytoplasm where it is converted back to pyruvate and lactate by pyruvate kinase (Pkm) and lactate dehydrogenase (Ldh) respectively. In our metabolomics analysis, we were unable to detect OAA and PEP. However, increase in lactate m+3, but not in m+2, and the accompanied decline in the labeling of TCA intermediates like fumarate m+2 and malate m+2, indicated reduction in TCA cycle activity and increased routing of pyruvate towards lactate in DR Paneth cells. In striking contrast, m+2 labeling of TCA cycle intermediates was increased in ISC under DR (Fig. 2E). Taken together, our metabolomic data indicated that Paneth cells catabolize alanine to preferably generate lactate. Stem cells possess very minimal expression of *Gpt2*, suggesting that the lactate m+3 found in ISCs might be derived from Paneth cells. Once Paneth cell derived lactate is transported to ISCs, it fuels their TCA cycle. Interestingly, also aspartate m+2 labeling was increased significantly in DR ISCs but reduced in Paneth cells (Fig. 2E). This indicates that malate-aspartate shuttle is active particularly in ISCs – suggesting that ISCs, with the support of their paneth cell niche, likely utilize this process to support cell proliferation ^21, 22^ during reduced nutrient availability.

### Lactate mediates the Gpt2 dependent effects on Paneth cell niche function

The metabolomic data suggested that Gpt2 mediated alanine utilization in Paneth cells can generate lactate as a transferable metabolic currency to be shared with stem cells during DR. Therefore, we next analyzed the functional relevance of exogenous lactate for the ISCs. To test this, we isolated crypts with ISCs and Paneth cells from AL or DR mice and assessed their capacity to form *de novo* crypts during organoid culture. Consistent with our earlier findings, Gpti blocked the DR induced crypt formation in organoids, but did not affect the crypt formation in AL organoids (Fig. S4A). Interestingly, lactate supplementation rescued the effects of Gpti in DR organoids, but had no effect on AL organoids. Moreover, lactate supplementation promoted organoid formation during Gpt inhibition only in co-cultures where Paneth cells were from DR mice (Fig. 3A). These results were also interesting as they demonstrated that lactate had negligible effect on AL ISCs unless they were exposed to DR Paneth cells. To test whether this was due to ongoing additional signals from the DR Paneth cells, or due to intrinsic changes in ISCs that were induced by the DR niche, we cultured DR ISCs alone under conditions where they can form organoids without Paneth cells. Demonstrating that alanine catabolism is compartmentalized to Paneth cells, alanine supplementation to ISCs had no effect (Fig. 3B). However, lactate indeed promoted organoid formation by ISCs from DR mice, but had no effect on AL ISCs (Fig. 3B and Fig. S4B), indicating that DR niche induces alterations in lactate use of the ISCs, and those changes can impact ISCs even after the niche signals are severed. Highlighting the role of reduced mTORC1 activity in Paneth cells as a modulator of niche interactions, lactate supplementation enhanced the supportive function of the Paneth cells isolated from Rapamycin-primed organoids with a genetic deletion of *Gpt2* – even if ISCs were from AL mice (Fig. 3C).

**Fig. 3:**
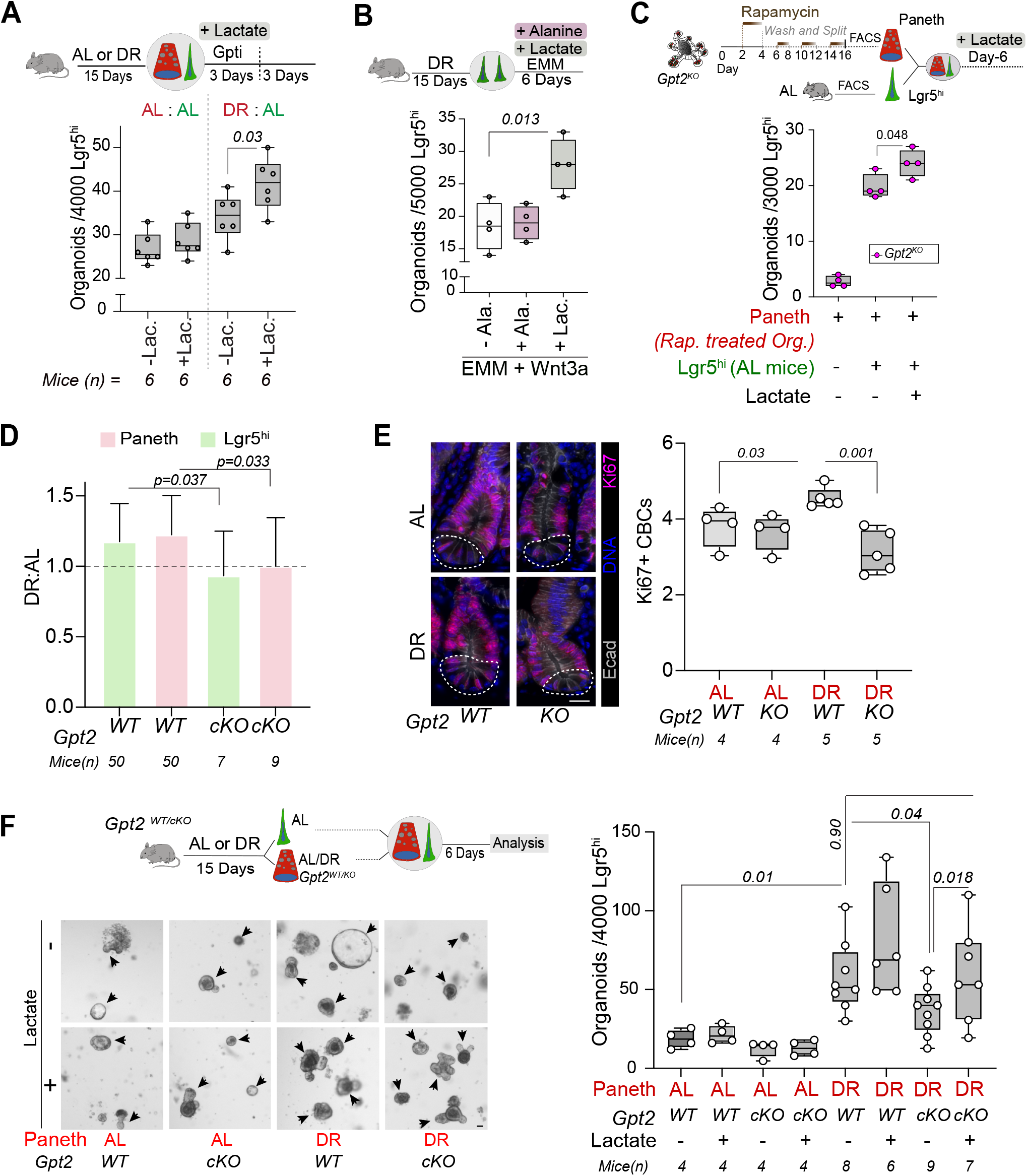
Lactate rescues the defective niche functions of *Gpt2*^*cKO*^ DR Paneth cells. **(A)** Lgr5^hi^ cells co-cultured with Paneth cells treated with Gpti (0.5 mM) and lactate (10 mM) for initial 3 days and organoid formation potential was scored on day 6 (n = 6 mice). **(B)** Clonogenic capacity of isolated Lgr5^hi^ cells from DR mice cultured in EMM media with addition of alanine and lactate (n = 4 mice). **(C)** Rapamycin treated *Gpt2*^*KO*^ Paneth cells were co-cultured with AL Lgr5^hi^ cells with and without lactate and clonogenic potential of Lgr5^hi^ cells was scored on day 6 (n=4 experiments). **(D)** Flow cytometric analysis of the Lgr5^hi^ and Paneth cells isolated from DR and AL *Gpt2*^*WT*^ and *Gpt2*^*cKO*^ mice. Population of Lgr5^hi^ and Paneth cells are shown as in the ratios of DR:AL (n=7-50 mice were analyzed per group). **(E)** Representative images of the immunohistochemical analysis of Ki67+ crypt base columnar cells (shown in the white dotted area), juxtaposed between the Paneth cells, in the small intestine of *Gpt2*^*WT*^ and *Gpt2*^*cKO*^ mice subjected to DR and AL (n=4-5 mice were analyzed). Ki67 (red), DNA (Blue) and E-cad (Gray). Scale bar 50 μm. **(F)** Representative micrographs show the clonogenic capacity of Lgr5^hi^ cells (isolated from *Gpt2*^*WT*^ AL mice) co-cultured with Paneth cells (with and without 10 mM lactate) isolated from AL and DR *Gpt2*^*WT*^ and *Gpt2*^*KO*^ mice (n=4-9 mice). Black arrows show the formed organoids at day 6. Two-tailed Student’s paired *t*-test was applied between the organoid cultures, where cells were isolated from the same mice but cultured with or without lactate. Scale bar 50 μm. Unless otherwise mentioned, data are shown as mean ± s.d. with two-tailed unpaired Student’s *t*-test. *p* values are shown in the corresponding panels. *p* <0.05 is considered statistically significant.

To further probe the role of Gpt2 *in vivo* in Paneth cells, we generated a mouse model harbouring a conditional allele of *Gpt2* (*Gpt2*^*fl*^), and crossed it with *Lgr5-CreERT2-IRES-EGFP* ^3^ to target Gpt2 in the intestinal epithelium (*Gpt2*^*cKO*^)(Fig. S5A). Since the Paneth cells can persist at the crypt base for months ^23^, we kept the mice for 90 days after 2 weeks of a tamoxifen supplemented diet to target *Gpt2* in Paneth cells (Fig. S5B-C). Upon 15 days of DR, *Gpt2*^*WT*^ and *Gpt2*^*cKO*^ conditional knock-out mice showed similar reduction of body weight (Fig. S5D). However, whereas DR increased the number of stem and Paneth cells in *Gpt2*^*WT*^ mice - as expected based on previous works ^2^ - DR did not induce changes in the cellular ratios in *Gpt2*^*cKO*^ mice (Fig. 3D). Moreover, the DR induced increase in the number of Ki67+ ISCs in crypts (recognized as crypt base columnar cells based on their location between Paneth cells) was lost in *Gpt2*^*cKO*^ mice (Fig. 3E).

We next probed the functional impact of *Gpt2* loss on intestinal regenerative capacity, and noted that the positive effects of DR on regenerative growth (Fig. S5E) were lost in crypts isolated from *Gpt2*^*cKO*^ mice. To address if these effects were indeed due to targeting of Gpt2 specifically in the Paneth cells, we isolated *Gpt2*^*WT*^ and *Gpt2*^*cKO*^ Paneth cells from AL and DR mice, and cultured them with *Gpt2*^*WT*^ ISC from AL mice. The DR induced boost in niche function was significantly reduced in *Gpt2*^*cKO*^ Paneth cells (Fig. 3F and S5F). Moreover, exogenous lactate restored the effects of DR in organoid cultures with *Gpt2*^*cKO*^ Paneth cells or crypts (Fig. 3F and S5E-F), indicating that the *Gpt2*^*cKO*^ Paneth cells were not defective otherwise and that lactate supplementation circumvents the effects of genetic Gpt2 targeting in the intestinal crypt.

Finally, to uncover how lactate is transferred from the Paneth cell niche to ISCs, we investigated the expression of putative lactate transporters ^24, 25^ and found that *Mct1/Slc16A1* is expressed at a higher level in ISCs than in the Paneth cells (Fig. 4A-B). To address the role of Mct1 in ISCs function, we used two potent inhibitors of Mct1: SR13800 ^26^, and AZD3965 ^27^. Both SR13800 and AZD3965 markedly blunted the regenerative growth of organoids from DR crypts but had only modest effects on AL crypts (Fig. S6A-B). Similarly, regenerative potential of crypts from Rapamycin treated mice was reduced by Mct1 inhibition (Fig. S6C). To test if these effects were due to Mct1 inhibition in ISCs, we exposed isolated ISCs to SR13800 before co-culture with untreated Paneth cells. While SR13800 did not impact organoid formation in co-cultures with AL Paneth cells, the boost by DR Paneth cells was significantly reduced (Fig. 4C). Importantly, the effect of SR13800 on organoid formation was restricted to co-cultures with DR Paneth cells, indicating that stem cells can form organoids normally under Mct1 inhibition, but only the additional metabolic support by DR Paneth cells is blocked. To test if this metabolic support takes effect via stem cell mitochondria, we blocked the mitochondrial pyruvate carrier with UK-5099 ^10^. Under conditions where DR stem cells are cultured without Paneth cells, the lactate mediated boost on organoid formation was significantly reduced upon UK-5099 treatment (Fig. 4D). Jointly, these results support the model where Paneth cell derived lactate is imported to ISCs during DR, and in ISCs lactate is converted to pyruvate to fuel the mitochondrial TCA cycle (Fig. 4E).

**Fig. 4:**
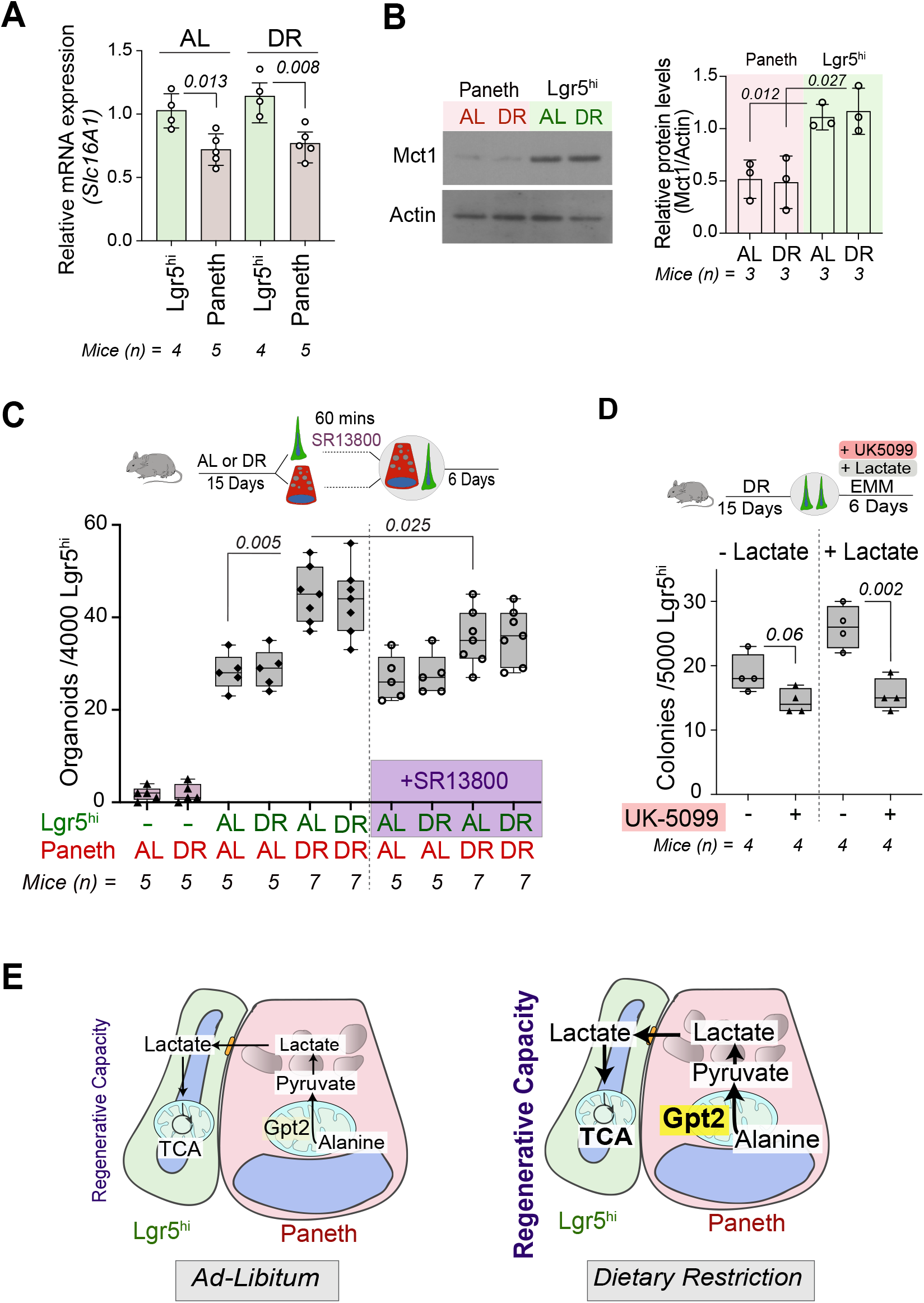
Lactate derived from Paneth cells fuels TCA cycle of stem cells under DR. **(A)** RT-qPCR of sorted Lgr5^hi^ and Paneth cells for *Slc16A1/Mct1* from AL and DR mice (n= 4-5 mice as indicated in figure). **(B)** Immunoblot and quantification of sorted Lgr5^hi^ and Paneth cells for Slc16A1/Mct1 from AL and DR mice (n= 3 mice) **(C)** Organoid formation potential of sorted Lgr5^hi^ cells from AL or DR mice treated with SR13800 (5 μM) for 60 mins and co-cultured with AL or DR Paneth cells was scored on day 6 (n =5-7 mice). **(D)** Clonogenic capacity of isolated Lgr5^hi^ cells from DR mice cultured in EMM media treated with UK-5099 (10μM) and the addition of lactate, was scored on the day-6 (n = 4 mice). **(E)** Schematic of the model for Paneth cell niche mediated metabolic crosstalk in governing the ISC function under DR. Unless otherwise mentioned, data are shown as mean ± s.d. and two-tailed unpaired student’s *t*-test was used. *p* values shown in the corresponding panels. *p* <0.05 is considered statistically significant.

## Discussion

Reduction in food intake promotes stem cell function and regeneration in multiple tissues ^2, 28–30^. However, underlying mechanisms that allow increase in stem cell functions under nutritional challenges have remained unknown. This conundrum is particularly apparent for the constantly and rapidly dividing stem cells of the intestine, where the source of energy and “building blocks” for the increase in ISC activity during DR ^2, 7^ has remained unknown. We discovered that during DR, Paneth cells of the stem cell niche rewire their own metabolism via Gpt2 to produce a transferrable metabolic currency, and thereby enable an increase in stem cell activity. The transferred metabolite, lactate, could not induce stem cell activity alone, highlighting the requirement of also other niche-associated signaling mechanisms ^2^ during DR.

Transferrable lactate was generated in Paneth cells via part of the gluconeogenic pathway, starting with catabolism of alanine by Gpt2. While expression of *Gpt2* in the *ad libitum* intestine is highest in Goblet and Paneth cells ^14^, *Gpt1* that can catalyze the same reaction is abundantly expressed by enterocytes. As enterocytes responsible for the nutrient absorption are located at the villus, whereas Paneth cells reside at the bottom of the crypts and are intermingled between ISCs, the Gpt2 dependent metabolic niche function is spatially separated from the Gpt1 transamination in the villus. Moreover, the DR-mediated induction of the Gpt2 but not Gpt1 was specific to crypts. Further studies separating these two routes of epithelial alanine catabolism are necessary to for example address their respective roles in the intestinal gluconeogenesis that has recently been found to play an important role in regulating systemic nutrient responses and counteract obesity and diabetes ^31–35^.

Muscle tissue and the intestine can produce and secrete alanine, which upon transport to the liver is used to drive gluconeogenesis that in turn allows the liver to either store or supply glucose to other organs ^36, 37^. This mechanism maintaining the systemic homeostasis of energy metabolism is known as Cahill cycle ^38^. Under prolonged fasting muscles will break down proteins and promote alanine secretion to the liver ^39^, whereas the utilization of alanine in the intestine under nutrient restriction remains understudied. In the light of the aforementioned compartmentalization of Gpt1 and Gpt2 driven alanine catabolism in the intestine and the possible inter-tissue exchange of metabolites, it will be interesting for future studies to address the source of alanine used by the Paneth cells during DR. Our metabolic tracing experiments demonstrated that intake of alanine by Paneth cells is increased during DR. Paneth cells have a particularly high expression of the neutral amino acid transporter *Slc1a4* that is responsible for alanine transport, and of the *Slc36a1* alanine lysosomal exporter ^11^. Moreover, inhibition of ketone body metabolism downregulates *Slc1a4* in Paneth cells ^40^, suggesting a possibility of unexplored bi-directional metabolic dialogue. Paneth cells may, therefore, harvest alanine from various intra- and extracellular sources depending for example on the natural fluctuations of food due to circadian rhythms.

The regulation of stem cells by growth and transcription factors has been extensively illuminated, but recent evidences also highlight the dynamic nature and broad involvement of metabolites in tissue adaptation and homeostatic control of stem cells ^2, 6, 7, 41, 42^. While exchange of metabolites between the niche and stem cells, and the importance of this exchange for mitochondrial metabolism in stem cells has previously been demonstrated ^8^, earlier work lacked real-time methods to evaluate the level of crosstalk. By establishing a novel metabolic tracing assay, we were able to determine that this mode of niche support is dynamically regulated and a necessary enabler for stem cell activity when nutrient availability is reduced. Notably, during this metabolic crosstalk, the Paneth cell niche catabolizes their alanine pool, however, the formed metabolites are not used for fueling its own metabolism, but instead shuttle to support the metabolism and proliferation of stem cells. Efflux of lactate from paneth niche may account for the expression of specific lactate exporters (MCTs). We also previously found that the central integrator of cues on nutrient status, mTORC1, mediates an increase in Paneth cell niche function by increasing generation of cADPR during DR and Rapamycin treatment ^2^. Recapitulating the DR effect by using mTOR inhibitor Rapamycin *in-vitro*, we found that the Gpt2 mediated Alanine catabolism of Paneth cells could modulate the lactate exchange between Paneth and stem cells and ensure the enhance stem cell activity possibly by fueling the TCA cycle. As DR reduces mTORC1 activity and protein synthesis ^43^, more Alanine might become available for lactate production during DR. Consequently, the two Paneth cell mechanisms, Gpt2 and cADPR, that both increase stem cell function during DR, may indeed be linked on the level of mTORC1 but appear to affect stem cells via separate mechanisms. As cADPR can stimulate the function of ISCs from AL mice while lactate cannot, but effects of cADPR are dependent on Paneth cell alanine metabolism by Gpt2, cADPR may provide a signal to ISCs on reduction in resources and evoke the ability of ISCs to import and utilize the niche derived lactate. Jointly these mechanisms adapt intestinal function to changing availability of food by building up regenerative capacity instead of differentiated cells when food is scarce and long absorptive villi are unnecessary. The insights from these findings may provide a scope for dietary and therapeutic interventions to promote intestinal regeneration during aging or to target pathologies.

## Acknowledgements

We thank J. Bärlund and M. Simula for the technical assistance. We also thank E. Kuuluvainen for the technical assistance in analysis of metabolomics data. We highly appreciate the services of Laboratory Animal Center, University of Helsinki.

## Notes

**Completing interests** Authors declare they have no conflict of interests.

**Grant support** PK and his laboratory members were supported by the Academy of Finland (266869, 304591 and 320185), ERC CoG 101045009, Swedish Research Council (2018-03078, 2022-01304), Cancerfonden 190634, the Jane and Aatos Erkko Foundation and the Cancer Foundation Finland. SI was supported by Magnus Ehrnrooth Foundation.

### Competing Interest Statement

The authors have declared no competing interest.

## References

1. Laplante M, Sabatini DM. mTOR signaling in growth control and disease. Cell 2012;149:274–93.

2. Yilmaz OH, Katajisto P, Lamming DW, et al. mTORC1 in the Paneth cell niche couples intestinal stem-cell function to calorie intake. Nature 2012;486:490–5.

3. Barker N, van Es JH, Kuipers J, et al. Identification of stem cells in small intestine and colon by marker gene Lgr5. Nature 2007;449:1003–7.

4. McCarthy N, Kraiczy J, Shivdasani RA. Cellular and molecular architecture of the intestinal stem cell niche. Nat Cell Biol 2020;22:1033–1041.

5. Cheng CW, Biton M, Haber AL, et al. Ketone Body Signaling Mediates Intestinal Stem Cell Homeostasis and Adaptation to Diet. Cell 2019;178:1115–1131 e15.

6. Mihaylova MM, Cheng CW, Cao AQ, et al. Fasting Activates Fatty Acid Oxidation to Enhance Intestinal Stem Cell Function during Homeostasis and Aging. Cell Stem Cell 2018;22:769–778 e4.

7. Igarashi M, Guarente L. mTORC1 and SIRT1 Cooperate to Foster Expansion of Gut Adult Stem Cells during Calorie Restriction. Cell 2016;166:436–450.

8. Rodriguez-Colman MJ, Schewe M, Meerlo M, et al. Interplay between metabolic identities in the intestinal crypt supports stem cell function. Nature 2017;543:424–427.

9. Sousa CM, Biancur DE, Wang X, et al. Pancreatic stellate cells support tumour metabolism through autophagic alanine secretion. Nature 2016;536:479–83.

10. Schell JC, Wisidagama DR, Bensard C, et al. Control of intestinal stem cell function and proliferation by mitochondrial pyruvate metabolism. Nat Cell Biol 2017;19:1027–1036.

11. Pentinmikko N, Iqbal S, Mana M, et al. Notum produced by Paneth cells attenuates regeneration of aged intestinal epithelium. Nature 2019;571:398–402.

12. Dohla J, Kuuluvainen E, Gebert N, et al. Metabolic determination of cell fate through selective inheritance of mitochondria. Nat Cell Biol 2022;24:148–154.

13. Van Lidth de Jeude JF, Vermeulen JL, Montenegro-Miranda PS, et al. A protocol for lentiviral transduction and downstream analysis of intestinal organoids. J Vis Exp 2015.

14. Haber AL, Biton M, Rogel N, et al. A single-cell survey of the small intestinal epithelium. Nature 2017;551:333–339.

15. Ouyang Q, Nakayama T, Baytas O, et al. Mutations in mitochondrial enzyme GPT2 cause metabolic dysfunction and neurological disease with developmental and progressive features. Proc Natl Acad Sci U S A 2016;113:E5598–607.

16. Ron-Harel N, Ghergurovich JM, Notarangelo G, et al. T Cell Activation Depends on Extracellular Alanine. Cell Rep 2019;28:3011–3021 e4.

17. Beuster G, Zarse K, Kaleta C, et al. Inhibition of alanine aminotransferase in silico and in vivo promotes mitochondrial metabolism to impair malignant growth. J Biol Chem 2011;286:22323–30.

18. Gray LR, Sultana MR, Rauckhorst AJ, et al. Hepatic Mitochondrial Pyruvate Carrier 1 Is Required for Efficient Regulation of Gluconeogenesis and Whole-Body Glucose Homeostasis. Cell Metab 2015;22:669–81.

19. Sebastian C, Ferrer C, Serra M, et al. A non-dividing cell population with high pyruvate dehydrogenase kinase activity regulates metabolic heterogeneity and tumorigenesis in the intestine. Nat Commun 2022;13:1503.

20. Hagopian K, Ramsey JJ, Weindruch R. Caloric restriction increases gluconeogenic and transaminase enzyme activities in mouse liver. Exp Gerontol 2003;38:267–78.

21. Garcia-Bermudez J, Baudrier L, La K, et al. Aspartate is a limiting metabolite for cancer cell proliferation under hypoxia and in tumours. Nat Cell Biol 2018;20:775–781.

22. Rabinovich S, Adler L, Yizhak K, et al. Diversion of aspartate in ASS1-deficient tumours fosters de novo pyrimidine synthesis. Nature 2015;527:379–383.

23. Roth S, Franken P, Sacchetti A, et al. Paneth cells in intestinal homeostasis and tissue injury. PLoS One 2012;7:e38965.

24. Sandforth L, Ammar N, Dinges LA, et al. Impact of the Monocarboxylate Transporter-1 (MCT1)-Mediated Cellular Import of Lactate on Stemness Properties of Human Pancreatic Adenocarcinoma Cells dagger. Cancers (Basel) 2020;12.

25. Bosshart PD, Kalbermatter D, Bonetti S, et al. Mechanistic basis of L-lactate transport in the SLC16 solute carrier family. Nat Commun 2019;10:2649.

26. Khan A, Valli E, Lam H, et al. Targeting metabolic activity in high-risk neuroblastoma through Monocarboxylate Transporter 1 (MCT1) inhibition. Oncogene 2020;39:3555–3570.

27. Beloueche-Babari M, Casals Galobart T, Delgado-Goni T, et al. Monocarboxylate transporter 1 blockade with AZD3965 inhibits lipid biosynthesis and increases tumour immune cell infiltration. Br J Cancer 2020;122:895–903.

28. Cerletti M, Jang YC, Finley LW, et al. Short-term calorie restriction enhances skeletal muscle stem cell function. Cell Stem Cell 2012;10:515–9.

29. Chen J, Astle CM, Harrison DE. Hematopoietic senescence is postponed and hematopoietic stem cell function is enhanced by dietary restriction. Exp Hematol 2003;31:1097–103.

30. Forni MF, Peloggia J, Braga TT, et al. Caloric Restriction Promotes Structural and Metabolic Changes in the Skin. Cell Rep 2017;20:2678–2692.

31. De Vadder F, Kovatcheva-Datchary P, Goncalves D, et al. Microbiota-generated metabolites promote metabolic benefits via gut-brain neural circuits. Cell 2014;156:84–96.

32. Soty M, Gautier-Stein A, Rajas F, et al. Gut-Brain Glucose Signaling in Energy Homeostasis. Cell Metab 2017;25:1231–1242.

33. Sun D, Wang K, Yan Z, et al. Duodenal-jejunal bypass surgery up-regulates the expression of the hepatic insulin signaling proteins and the key regulatory enzymes of intestinal gluconeogenesis in diabetic Goto-Kakizaki rats. Obes Surg 2013;23:1734–42.

34. Troy S, Soty M, Ribeiro L, et al. Intestinal gluconeogenesis is a key factor for early metabolic changes after gastric bypass but not after gastric lap-band in mice. Cell Metab 2008;8:201–11.

35. Vily-Petit J, Soty-Roca M, Silva M, et al. Intestinal gluconeogenesis prevents obesity-linked liver steatosis and non-alcoholic fatty liver disease. Gut 2020;69:2193–2202.

36. Felig P. The glucose-alanine cycle. Metabolism 1973;22:179–207.

37. Potts A, Uchida A, Deja S, et al. Cytosolic phosphoenolpyruvate carboxykinase as a cataplerotic pathway in the small intestine. Am J Physiol Gastrointest Liver Physiol 2018;315:G249–G258.

38. Sarabhai T, Roden M. Hungry for your alanine: when liver depends on muscle proteolysis. J Clin Invest 2019;129:4563–4566.

39. Pozefsky T, Tancredi RG, Moxley RT, et al. Effects of brief starvation on muscle amino acid metabolism in nonobese man. J Clin Invest 1976;57:444–9.

40. Gebert N, Cheng CW, Kirkpatrick JM, et al. Region-Specific Proteome Changes of the Intestinal Epithelium during Aging and Dietary Restriction. Cell Rep 2020;31:107565.

41. Beyaz S, Mana MD, Roper J, et al. High-fat diet enhances stemness and tumorigenicity of intestinal progenitors. Nature 2016;531:53–8.

42. Peregrina K, Houston M, Daroqui C, et al. Vitamin D is a determinant of mouse intestinal Lgr5 stem cell functions. Carcinogenesis 2015;36:25–31.

43. Saxton RA, Sabatini DM. mTOR Signaling in Growth, Metabolism, and Disease. Cell 2017;168:960–976.

