## Supplementary material for "Alanine catabolism of Paneth cells maintains intestinal stem cell function during dietary restriction": Figure legends

**Supplementary Figure legends:**

**Fig. S1: Gpt2 is induced upon DR or DR-mimetic Rapamycin treatment.**
**(A)** Immunoblotting of isolated crypts (consists of many other cell types than Paneth cells and $Lgr5^{hi}$ stem cells) from AL and DR mice for Gpt2 suggests a modest increase upon DR. Densitometric quantification (ratio to actin) (n = 3 mice per group). **(B)** Immunohistochemical (IHC) analysis of intestinal stem cells (Olfm4) and Paneth cells (Lysozyme 1) in AL and DR mice. Activation of mTOR was assessed by ph-S6 staining (n=5 mice). Scale Bar 20μm. **(C)** Crypts and villus cell lysates were prepared from AL and DR mice. Represented images are indicating the purity index. Immunoblotting was done to analyze the levels of Gpt1 and Gpt2 (n=3 mice). **(D)** Knockout efficacy of *Gpt2* in organoids was verified by RT-PCR and immunoblotting. *Rpl13a* was used as an internal control. Regenerative images of the *Gpt2* knockout (KO) organoids. Scale bar 50μm. **(E)** Immunoblotting and quantification of Rapamycin primed scrambled and $Gpt2^{KO}$ organoids indicates a modest increase in Gpt2 levels on rapamycin priming (n=3 mice). **(F)** $Lgr5^{hi}$ cells co-cultured with Paneth cells isolated from AL mice treated with Gpti and 2 nM Rapamycin for the initial 3 days. Clonogenic potential of stem cells was calculated on day 6 (n=4 mice). Scale bar 50μm. Unless otherwise mentioned, data are shown as mean ± s.d.; two-tailed unpaired Student’s *t*-test. *p* values shown in the corresponding panels. $p<0.05$ is considered statistically significant.

**Fig. S2 Blockade of alanine metabolism reduces ISC function under DR.**
**(A)** $Lgr5^{hi}$ cells co-cultured with Paneth cells treated with Gpti (0.5mM) and cyclic-ADPR (50μM) for initial 3 days and organoid formation potential was scored on 6 day. (n = 5 mice), scale bar 50 μm. **(B)** $Lgr5^{hi}$ cells co-cultured with Paneth cells treated with Gpti and cyclic-ADPR for 6 days and organoid formation potential was scored on day 6. (n = 4 mice). Crypt budding potential of formed organoids from the co-culture of DR Paneth cells and AL $Lgr5^{hi}$ with initial (3 days) of Gpti was compared (n = 4 mice). **(C)** Composition of the ENR mimetic media (EMM). EMM lacking alanine was formulated based on MEM. **(D)** Organoids derived from AL or DR mice using ENR and EMM culture medium. Alanine (0.5mM) and Gpti (0.5mM) were added and the generation of *de novo* crypt domains per organoid was scored on day 5-6 (n= 4-5 mice). Scale bar 50 μm **(E)** Isolated crypts from AL and DR mice was cultured in EMM with and without alanine supplementation and organoid forming capacity (starting frequency) of crypts was assessed (day 2) (n = 5 mice). Unless otherwise mentioned, data are shown as mean ± s.d. with two-tailed unpaired Student’s *t*-test; *P* values shown in the corresponding panels. $P<0.05$ is considered statistically significant.

**Fig. S3. Alanine catabolism in DR Paneth cells forms lactate that supports TCA cycle in ISCs.**

**(A)** Percentage pool of labeled m+3 alanine and m+3 lactate to the total (m+0, m+1, m+2, m+3) pool of metabolites in $Lgr5^{hi}$ and Paneth cells in AL and DR state (n=4 mice). Data are presented as mean ± s.d. Two-tailed unpaired Student’s *t*-test was used. *p* values are shown in the corresponding panels. *P* <0.05 is considered statistically significant.

**Fig. S4. Lactate generation by DR Paneth cells regulates the ISC function.**
**(A)** Regenerative capacity of isolated crypts from AL or DR mice is shown as *de novo* crypts per organoid at day 5, with and without supplementation of Gpt inhibitor (0.5 mM) and lactate (10 mM). n=8-11 mice; scale bar 50µm. (**B**) Clonogenic capacity of isolated Lgr5$^{hi}$ cells from AL mice cultured in EMM media with addition of alanine and lactate was scored on the day 6. Unless otherwise mentioned, data are shown as mean ± s.d. Two-tailed unpaired Student's *t*-test was used. *p* values are shown in the corresponding panels. *p* <0.05 is considered significant.

**Fig. S5: Conditional deletion of *Gpt2* leads to reduced epithelial regeneration under DR in a lactate-dependent manner.** (**A**) Schematic for generating *Lgr5-CreERT2;Gpt2* $^{fl/fl}$ mice by targeting exon 4 of *Gpt2* (*Gpt2*$^{fl/fl}$). (**B**) Flow chart for tamoxifen (TAM) feeding and AL/DR dietary regimes of the mice. (**C**) qPCR data of the relative expression level of *Gpt2* in Paneth cells isolated from *Gpt2*$^{WT}$ and *Gpt2*$^{KO}$ DR mice (n=5-8 mice). *Rpl13a* was used as internal control. (**D**) Changes of the body weights of the mice after 15 days of dietary restriction. (**E**) Representative images of the organoids cultured with and without lactate (10 mM). Crypts were isolated from *Gpt2*$^{WT}$ and *Gpt2*$^{KO}$ mice under AL and DR (left). Box plot shows the *de novo* crypt formation in the organoids. **(F)** Representative micrographs show the crypt bud formation from single Lgr5$^{hi}$ cells. Regenerative capacity (crypt bud/organoid) of Lgr5$^{hi}$ cells (isolated from AL mice) co-cultured with Paneth cells ((with and without 10 mM lactate) isolated from AL and DR mice (n=4-5 mice) of *Gpt2*$^{WT}$ and *Gpt2*$^{KO}$ background. Brown arrows show the *de novo* crypt bud formation. Quantification was done on day 10. Scale bar 50 µm. Data are presented as mean ± s.d. Two-tailed unpaired Student's *t*-test was used. *p* values are shown in the corresponding panels. *p* <0.05 is considered statistically significant. Scale bar 50 µm.

**Fig. S6: Inhibition of Mct1 impairs epithelial regeneration under** DR. (**A**) Regenerative capacity of isolated crypts from AL or DR mice to form *de novo* crypts in formed organoid at day 5 treated with Mct1 inhibitor, SR13800 and (**B**) AZD3965. n=4 mice. (**C**) Regenerative capacity of isolated crypts from Rapamycin treated mice to form *de novo* crypts in formed organoid at day 6 treated with 500 nM Mct1 inhibitor, SR13800 (n=4 mice). Unless otherwise mentioned, data are shown as mean ± s.d. Two-tailed unpaired Student's *t*-test was used. *p* values are shown in the corresponding panels. *p* <0.05 is considered significant. Scale bar 50 µm.
