## Supplementary figures and images for "Alanine catabolism of Paneth cells maintains intestinal stem cell function during dietary restriction"

### Supplementary Figure 1

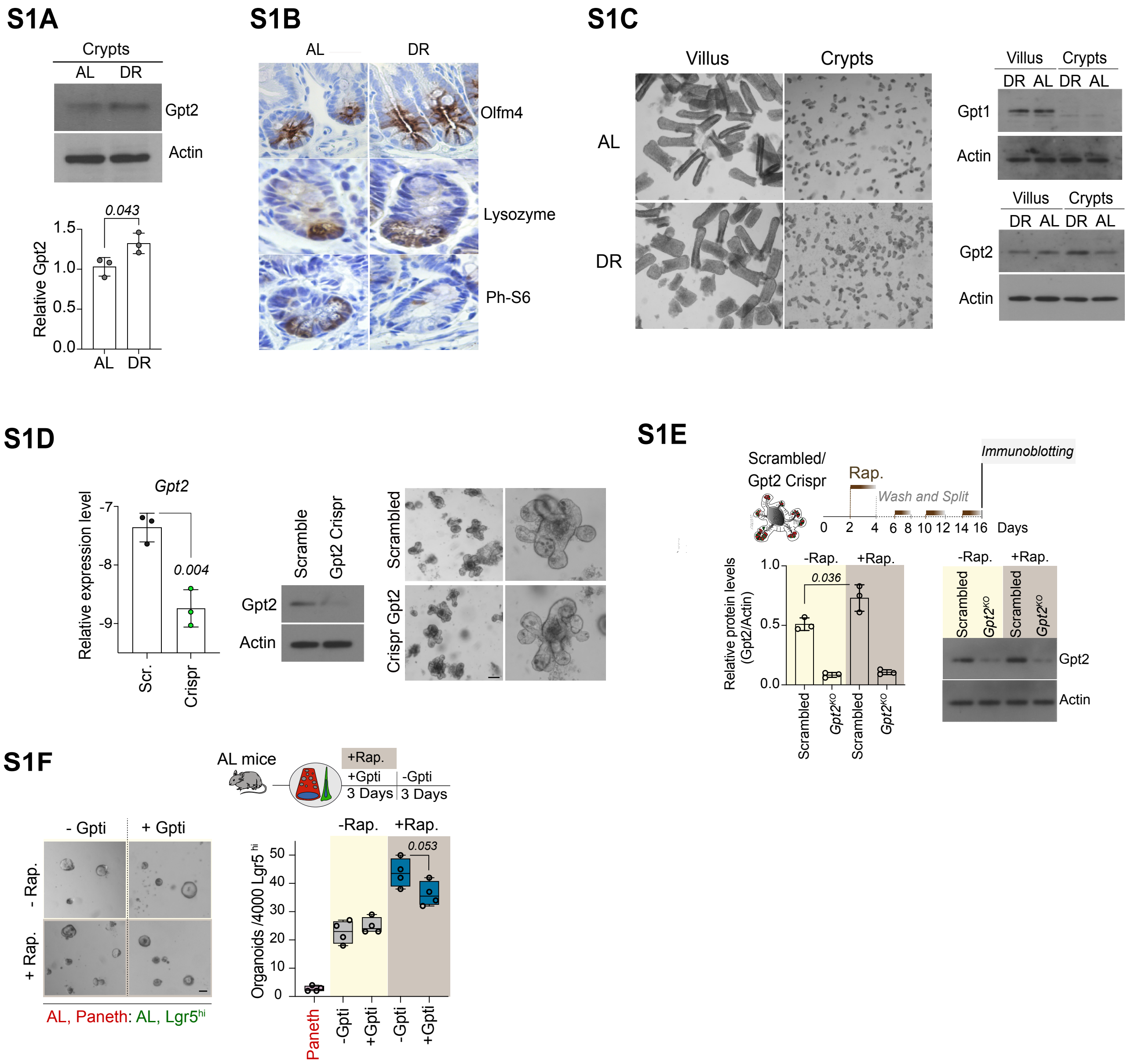


**Figure S1: Gpt2 is induced upon DR or DR-mimetic Rapamycin treatment.**

### Supplementary Figure 2

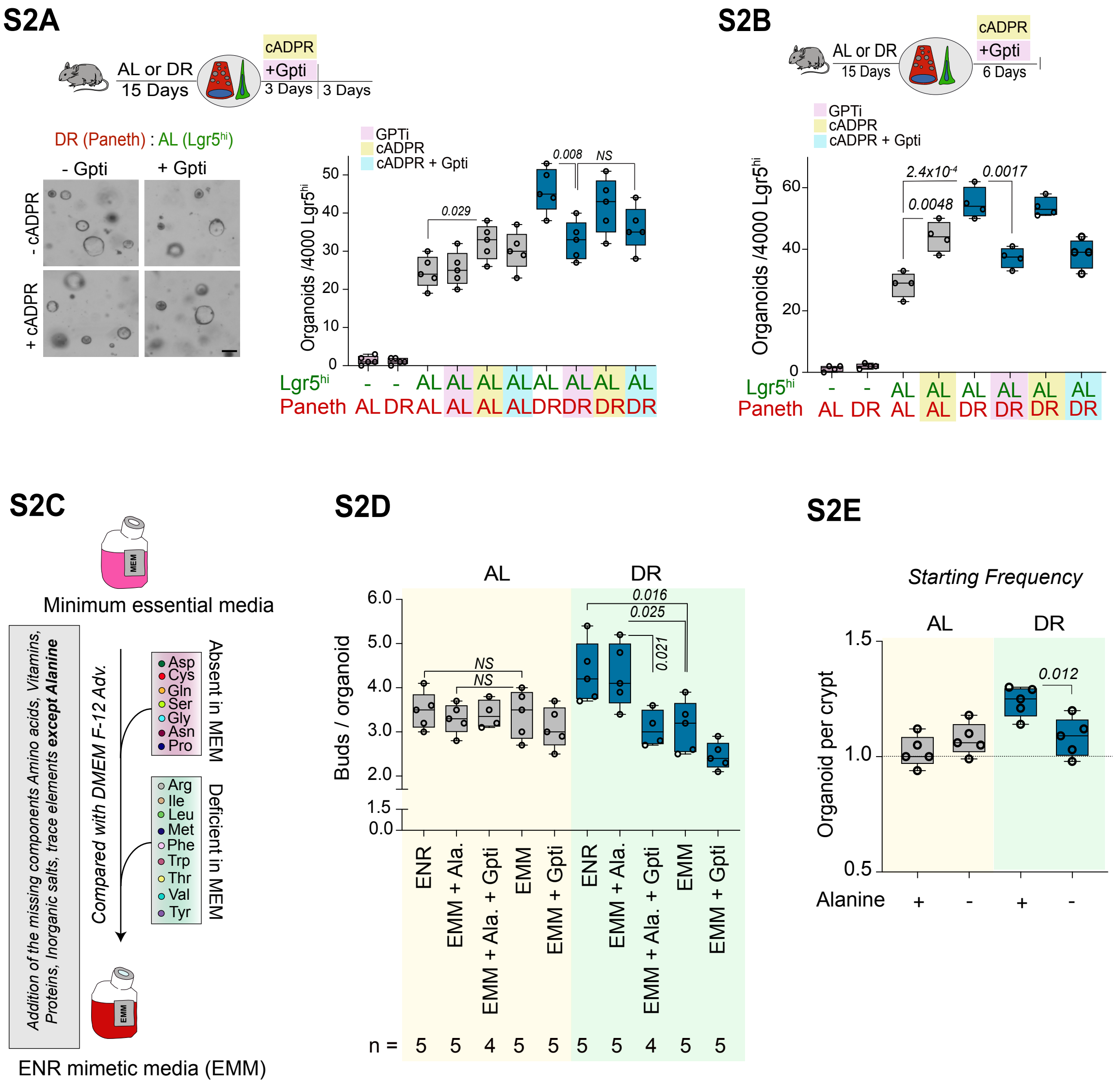


**Figure S2: Blockade of alanine metabolism decline stem cell function under DR.**

### Supplementary Figure 3

## S3A

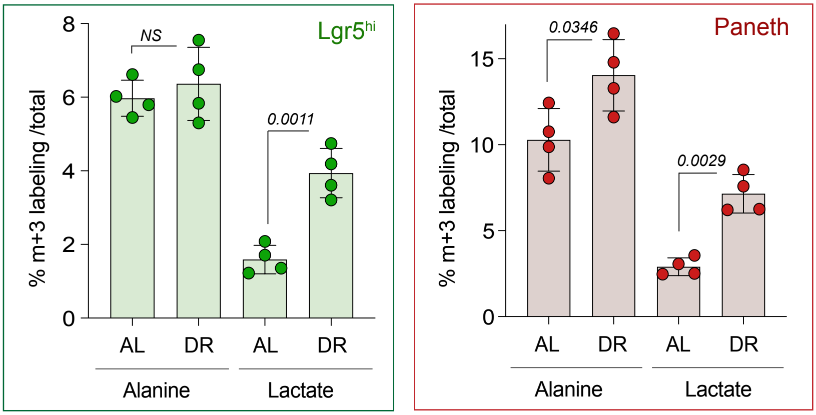


**Figure S3: Alanine catabolism in DR Paneth cells forms lactate that supports TCA cycle in ISCs.**

### Supplementary Figure 4

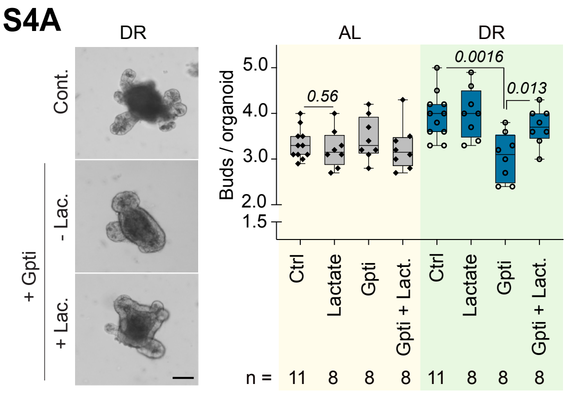


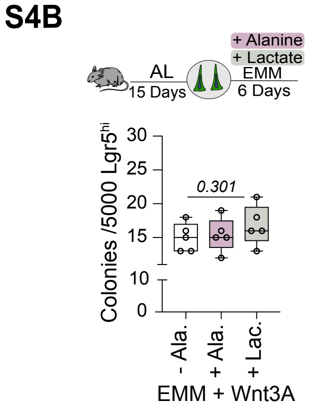


**Figure S4: Gpt inhibition counteracts the effects of lactate.**

### Supplementary Figure 6

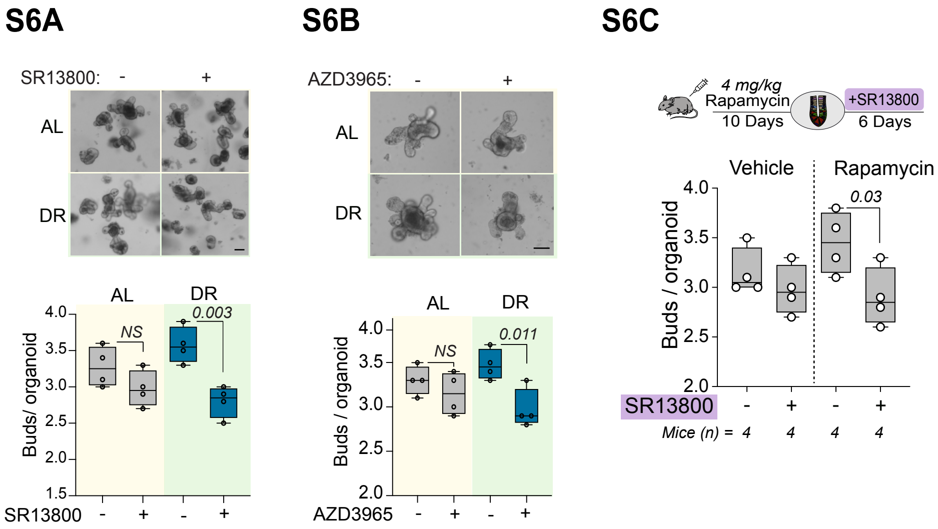


**Figure S6: Inhibition of Mct1 impairs epithelial regeneration under DR.**
