## Supplementary Figure 5 for "Alanine catabolism of Paneth cells maintains intestinal stem cell function during dietary restriction"

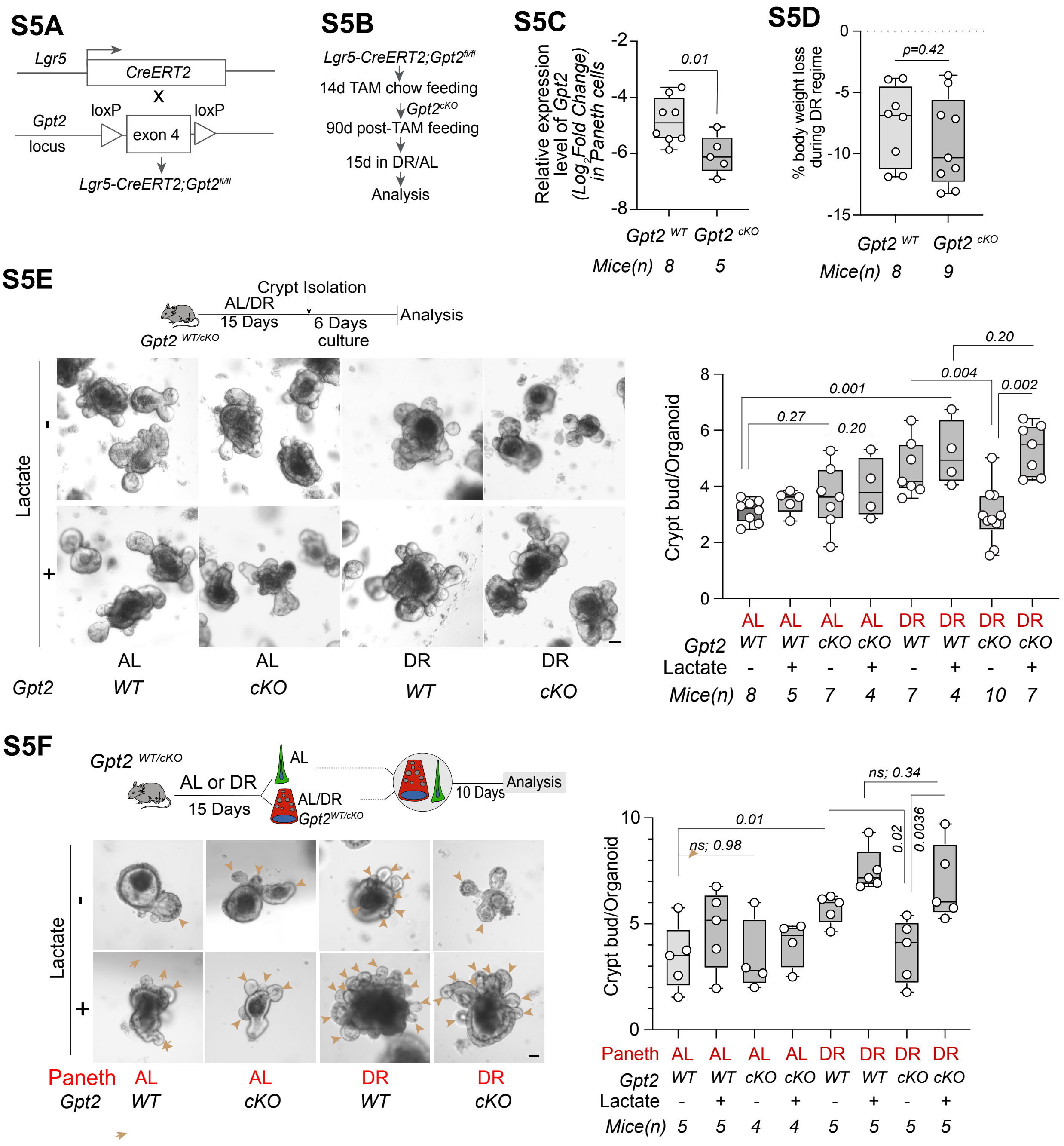


**Figure S5: Conditional deletion of *Gpt2* leads to reduced epithelial regeneration under DR in a lactate-dependent manner.**
